# Biosynthetic Enzyme-guided Disease Correlation Connects Gut Microbial Metabolites Sulfonolipids to Inflammatory Bowel Disease Involving TLR4 Signaling

**DOI:** 10.1101/2023.03.16.533047

**Authors:** Ethan A. Older, Jian Zhang, Zachary E. Ferris, Dan Xue, Zheng Zhong, Mary K. Mitchell, Michael Madden, Yuzhen Wang, Hexin Chen, Prakash Nagarkatti, Mitzi Nagarkatti, Daping Fan, Melissa Ellermann, Yong-Xin Li, Jie Li

**Affiliations:** Department of Chemistry and Biochemistry, University of South Carolina, Columbia, South Carolina, USA; Department of Chemistry and The Swire Institute of Marine Science, The University of Hong Kong, Pokfulam Road, Hong Kong, China; Southern Marine Science and Engineering Guangdong Laboratory (Guangzhou), Guangzhou, China; Department of Biological Sciences, University of South Carolina, Columbia, South Carolina 29209, United States; Department of Cell Biology and Anatomy, School of Medicine, University of South Carolina, Columbia, South Carolina 29209, United States; Department of Pathology, Microbiology and Immunology, School of Medicine, University of South Carolina, Columbia, South Carolina 29209, United States

## Abstract

The trillions of microorganisms inhabiting the human gut are intricately linked to human health. At the species abundance level, correlational studies have connected specific bacterial taxa to various diseases. While the abundances of these bacteria in the gut serve as good indicators for disease progression, understanding the functional metabolites they produce is critical to decipher how these microbes influence human health. Here, we report a unique biosynthetic enzyme-guided disease correlation approach to uncover microbial functional metabolites as potential molecular mechanisms in human health. We directly connect the expression of gut microbial sulfonolipid (SoL) biosynthetic enzymes to inflammatory bowel disease (IBD) in patients, revealing a negative correlation. This correlation is then corroborated by targeted metabolomics, identifying that SoLs abundance is significantly decreased in IBD patient samples. We experimentally validate our analysis in a mouse model of IBD, showing that SoLs production is indeed decreased while inflammatory markers are increased in diseased mice. In support of this connection, we apply bioactive molecular networking to show that SoLs consistently contribute to the immunoregulatory activity of SoL-producing human microbes. We further reveal that sulfobacins A and B, two representative SoLs, primarily target Toll-like receptor 4 (TLR4) to mediate immunomodulatory activity through blocking TLR4’s natural ligand lipopolysaccharide (LPS) binding to myeloid differentiation factor 2, leading to significant suppression of LPS-induced inflammation and macrophage M1 polarization. Together, these results suggest that SoLs mediate a protective effect against IBD through TLR4 signaling and showcase a widely applicable biosynthetic enzyme-guided disease correlation approach to directly link the biosynthesis of gut microbial functional metabolites to human health.

## Introduction

The human gut microbiome, composed of trillions of microorganisms, is increasingly recognized as having a significant influence on human health through the production of numerous functional metabolites that are in direct contact and constant exchange with host cells^1^. Metabolism at the human-microbiota interface has been formed and regulated by the long-term symbiosis and co-evolution between humans and microbes, inherently linking human microbial functional metabolites to human health^2–7^. At the species abundance level, numerous human microbes have been correlated with disease phenotypes, however, the functional metabolites which underlie these conditions remain largely unknown^8–10^. To expand our understanding of the complex host-microbe interactions that influence human health, human microbiome research has begun advancing to the next level of revealing the microbial functional metabolites and their corresponding molecular mechanisms that drive specific disease phenotypes^11, 12^. A remarkable class of such functional metabolites is microbe-derived lipids. Biosynthesized *de novo* by human gut microbiota or biotransformed from dietary nutrients, these lipids can be directly sensed by the host to modulate both metabolic and immunological pathways^13^. Many studies have focused on common microbe-derived lipids, such as short chain fatty acids (SCFAs) and phospholipids (PLs)^14, 15^, however, there are a significant number of underexplored lipids which are equally capable of influencing human health^16^.

In our search for such underexplored microbial lipids, our attention was drawn to *Alistipes* and *Odoribacter* as two commensal genera of the human gut microbiome with emerging significance in modulating health, particularly in the two primary forms of inflammatory bowel disease (IBD): Ulcerative colitis (UC) and Crohn’s disease (CD)^17–20^. Species abundances from both genera have been found to negatively correlate with IBD pathogenesis, suggesting that they may play a protective role against IBD^17, 18^. Additionally, some species have been shown to ameliorate symptoms of IBD^19, 20^. Notably, both *Alistipes* and *Odoribacter* are prolific sulfonolipid (SoL) producers^21^. SoLs are unique lipid molecules that bear striking structural similarity to both bacterial and endogenous sphingolipids (SLs), which are known for their role in mediating immune signaling in the human body^22^. Species of *Alistipes* and *Odoribacter* do not produce SLs^21, 23^, but instead produce SoLs in high abundance^21, 24^. The replacement of SLs by structurally related SoLs in *Alistipes* and *Odoribacter* suggests that SoLs may serve as functional metabolites and exert similar but distinctly different functions compared to their SL cousins. In fact, we have found that sulfobacin A (SoL A), a representative of human microbial SoLs produced by *Chryseobacterium gleum* F93 DSM 16776, exhibits unusual dual immunomodulatory activity *in vitro* by modulating inflammatory cytokine production, especially through suppression of the lipopolysaccharide (LPS)-induced inflammatory response^25^ which has been reported as a key contributor to the progression of IBD^26–29^. Whether SoLs produced by *Alistipes* and *Odoribacter*, two genera negatively associated with IBD^17, 18^, represent functional metabolites in this negative correlation is unknown. Furthermore, the molecular target(s) of SoLs as a whole class of unique and abundant lipids is also unknown.

To investigate the potential mechanistic role of SoLs as functional metabolites in the IBD-protective effects, we developed a biosynthetic enzyme-guided disease correlation approach which connects the biosynthesis of functional metabolites directly to human health conditions. While untargeted metabolomic methods have been developed to probe potential functional metabolite trends in disease, these methods face challenges such as the complexity of the metabolome, the lack of reference databases for identification, trace metabolite amounts, and a high degree of variability between different metabolomes, all of which lead to difficulty in revealing specific gut microbial functional metabolites as drivers of molecular mechanisms in disease. Owing to the widespread availability of large high-quality disease-related sequencing datasets, our unique approach takes advantage of specific critical biosynthetic enzymes required for the biosynthesis of corresponding functional metabolites. Increased or decreased expression of these enzymes, reflecting the production of a specific functional metabolite, can be correlated with different disease parameters to reveal positive or negative associations, which subsequently enables a more focused, targeted metabolomic analysis to rapidly filter out metabolites of interest and further confirm their association with disease. We apply our approach to gut microbiome sequencing data collected from real world IBD patients to link the abundance and expression of SoL-specific biosynthetic enzymes directly to IBD pathogenesis. Different from correlating microbial species abundance with disease followed by various efforts to identify functional metabolites, our approach streamlines this process in a targeted manner by analyzing the changes in their respective biosynthetic enzyme expression patterns in response to disease.

In this work, we revealed a negative correlation between SoL biosynthesis and human IBD pathogenesis through a unique biosynthetic enzyme-guided disease correlation approach followed by targeted chemoinformatic analysis. We then experimentally validated this informatic correlation using a mouse model of IBD. Through bioactive molecular networking, we determined that SoLs consistently contribute to the immunoregulatory activity of SoL-producing human gut commensals. Using cell-based assays, we also revealed that SoLs A and B primarily mediate their immunomodulatory activity through interaction with TLR4. Specifically, SoLs bind directly to TLR4 via the accessory protein myeloid differentiation factor 2 (MD-2) and displace LPS from MD-2 at higher concentrations, leading to suppression of TLR4 signaling pathways and macrophage M1 polarization. Together, our results demonstrate an informatics-based, functional metabolites-focused approach to uncovering the chemical basis and molecular mechanisms of intricate host-microbe interactions and outline a potential mechanism by which *Alistipes* and *Odoribacter*-derived SoLs exert a protective effect against IBD progression through disruption of LPS-mediated TLR4 signaling.

## Results

### Biosynthetic enzyme-guided disease correlation analysis reveals a negative correlation between SoLs biosynthesis and IBD

We began by systematically investigating the biosynthetic potential of SoLs from 285,835 human gut bacterial reference genomes including single amplified genomes (SAGs) and metagenome-assembled genomes (MAGs)^30^. Based on sequence homology with experimentally verified SoL biosynthetic enzymes^25, 31–33^ (**Supplementary Fig. 1, Supplementary Table 1**), we identified a total of 562,214 homologous enzyme sequences, including 469,012 cysteate synthases (CYS), 33,486 cysteate fatty acyltransferases (CFAT), and 59,716 short-chain dehydrogenases/reductases (SDR) (**Extended Data Fig. 1a**). Uncovering phylogenetic trends, we found that these three enzymes were widely distributed in 255,572 genomes across 21 phyla, with the majority belonging to Bacteroidota and Firmicutes_A (**Extended Data Fig. 1b**). A subset of 6.21% (15,863/255,572) of the genomes was found to encode all three putative SoL biosynthetic enzymes (**Extended Data Fig. 1a**). To prioritize them for further analysis, we filtered the homologs on the basis of three rules: (1) the homology of both CFAT and CYS must equal or exceed 50% sequence similarity with experimentally validated CFATs and CYSs (**Supplementary Table 1**), as these enzymes are the first two specific enzymes in the biosynthetic pathway of SoLs that distinguish the biosynthesis between SoLs and SLs^25, 31–33^; (2) the homologous regions of CYS, CFAT, and SDR should include protein domains with Pfam IDs PF00291, PF00155, and PF00106, respectively (hit score > 50); (3) a set of homologous enzymes, especially SDR enzymes that show variable sequence similarities, should come from the same genome encoding all three enzymes as all three are required for SoL biosynthesis, thus ensuring co-occurrence. Applying these rules, we prioritized 9,731 CYS (1,384 unique sequences), 9,740 CFAT (917 unique sequences), and 10,319 SDR enzymes (1,076 unique sequences) (**Fig. 1a**, **Supplementary Data 1**) from 9,633 bacterial genomes. The prioritized enzymes were distributed among 42 species from Bacteroidota and one species from Firmicutes_A (**Fig. 1b**, **Supplementary Data 2**). Of note, among the 42 species from Bacteroidota, 71% (30/42) of them belong to bacterial families that have been previously reported to produce SoLs (**Fig. 1b**) including *Rikenellaceae* (containing genera *Alistipes* and *Alistipes*_A)^21, 34^, *Marinifilaceae* (containing genus *Odoribacter*)^21^, and *Weeksellaceae* (containing genus *Chryseobacterium* B)^25, 35^.

**Fig. 1.**
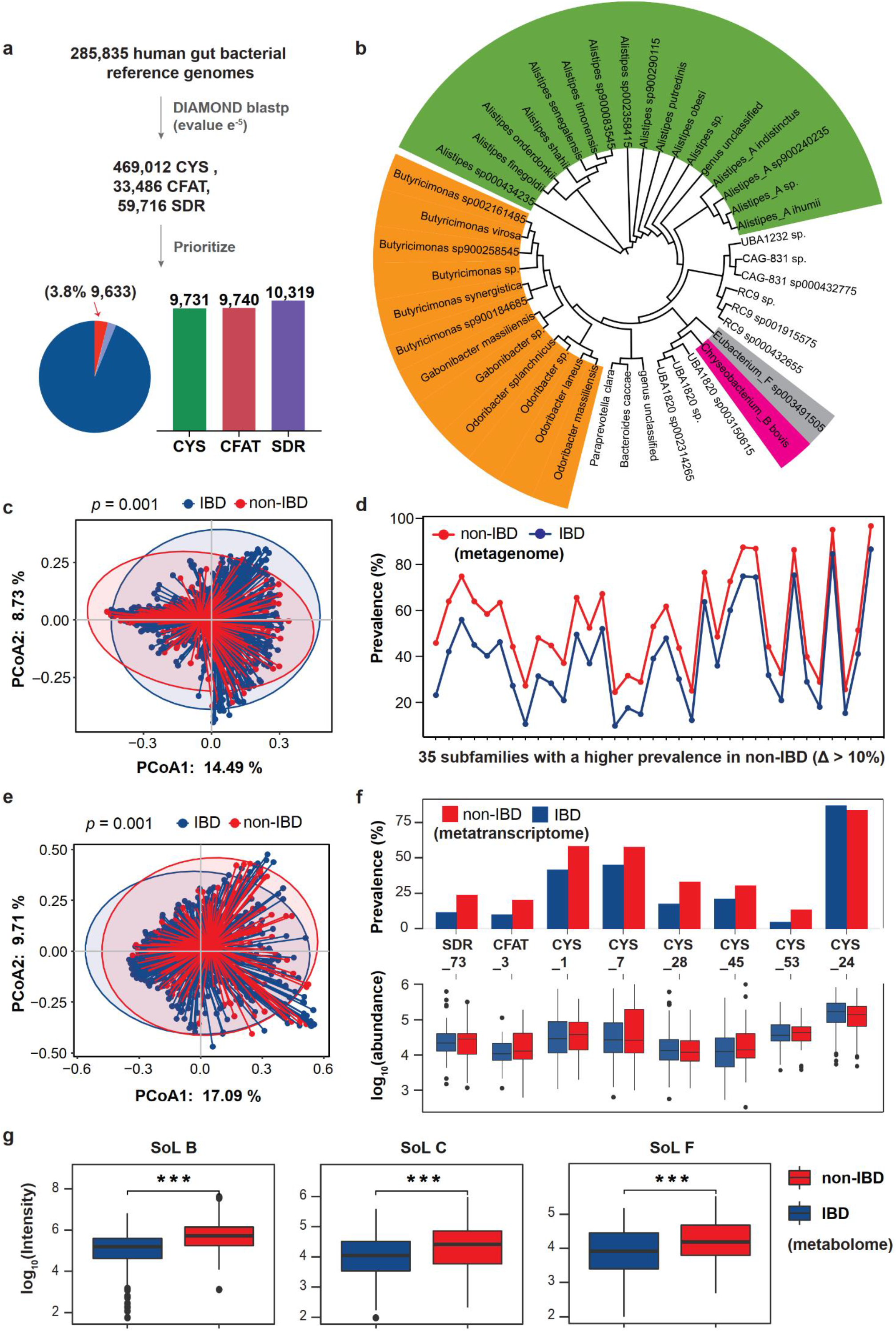
The presence and expression profiles of SoL biosynthetic enzymes and the production of SoLs differ in IBD subjects versus healthy controls. **a**, Overview of SoL biosynthetic enzymes identified in human gut bacteria. 562,214 putative SoL biosynthetic enzymes were identified across 21 bacterial phyla. 6.21% of genomes encode 3 types of SoL biosynthetic enzymes (Pie chart, sections in red and purple). Bar chart shows the number of prioritized SoL biosynthetic enzymes encoded by 9,633 genomes (highlighted in red in the pie chart). **b**, A circular phylogenetic tree shows the prioritized SoL biosynthetic enzymes found primarily in species from Bacteroidota (highlighted in green and orange). The tree is annotated with species names and colored by taxonomic families (*Rikenellaceae*: green; *Marinifilaceae*: orange; *Weeksellaceae*: pink; *Lachnospiraceae*: grey). **c**, Principal Coordinate Analysis (PCoA) shows differences in the presence profile of overall SoL biosynthetic enzyme subfamilies between IBD and non-IBD groups based on Jaccard distance. **d**, 35 SoL biosynthetic enzyme subfamilies were significantly more prevalent in healthy individuals (red dots) than in IBD groups (blue dots) (two-sided Fisher’s exact test, *p* < 0.05) with difference of prevalence > 10%. e, PCoA showing the differences in the expression profile of overall SoL biosynthetic enzyme subfamilies between IBD and non-IBD groups based on Bray-Curtis distances. **f**, Expression profiles of 8 differential SoL biosynthetic enzyme subfamilies (two-sided Mann-Whitney *U* test, adjusted *p* < 0.05). Upper panel: bar charts showing the prevalence of differential SoL biosynthetic enzyme subfamilies across non-IBD (red) and IBD individuals (dark blue). Statistical significance for prevalence was calculated using a two-sided Fisher’s exact test. Except CYS subfamily24 (no significance), all were significantly higher in prevalence in non-IBD than IBD groups (*p* < 0.05). Lower panel: box plots displaying the abundance profiles of differential SoL biosynthetic enzyme subfamilies in non-IBD (red) and IBD individuals (dark blue). Significance was further determined by one-sided Mann-Whitney *U* test, with adjusted *p*-value < 0.05. g, Box plots showing the absolute intensity of SoLs B, C, and F detected in non-IBD and IBD individuals collected from IBDMDB. Significance was determined by Wilcoxon rank sum test: * 0.01 < *p* < 0.05, ** 0.001 < *p* < 0 .01, and *** *p* < 0.001.

To determine whether there is a link between gut microbial capacity of producing SoLs and IBD incidence, we conducted a comparative analysis of metagenomic and metatranscriptomic data obtained from the Inflammatory Bowel Disease Multi’omics Database (IBDMDB, https://ibdmdb.org/)^18, 36^. We began by generating sequence similarity networks with a 90% sequence identity threshold to group enzymes with similar functions. Consequently, we categorized the prioritized biosynthetic enzymes into 214 subfamilies (79 CYS subfamilies; 25 CFAT subfamilies; and 110 SDR subfamilies) for the subsequent analyses (**Supplementary Table 2**). Looking for the presence of the prioritized 214 subfamilies in IBD cohorts, we identified 154 subfamilies in 667 metagenome samples (182 healthy samples and 485 IBD disease samples), of which 116 subfamilies were detected in ≥ 5% of samples (**Extended Data Fig. 1c**). Beta diversity of the presence of these 116 subfamily biosynthetic enzymes indicated that the overall composition of SoL biosynthetic enzyme subfamilies was significantly different between the healthy and IBD cohorts (**Fig. 1c**, Jaccard distance, PERMANOVA *p* = 0.001). Of note, 57 subfamilies had a significantly higher prevalence (Fisher’s exact test *p* < 0.05) in healthy individuals as compared to IBD cases (**Supplementary Table 3**), among which 35 subfamilies (18 CYS subfamilies, 2 CFAT subfamilies, and 15 SDR subfamilies) further show a difference of prevalence > 10% (**Fig. 1d**).

To further examine the difference between the expression profiles of SoL biosynthetic enzymes between the IBD and healthy groups, we extended our comparative analysis to the metatranscriptomic level. We found that 132 SoL biosynthetic enzyme subfamilies were expressed in 777 metatranscriptomic samples (193 healthy samples and 584 IBD disease samples), with about 42% (55/132) detected in at least 5% of samples (**Extended Data Fig. 1d**). Beta diversity of the expression profiles of SoL biosynthetic enzymes suggested that the overall expression of these enzyme subfamilies is significantly different between the healthy and IBD cohorts (**Fig. 1e**, Bray-Curtis distance, PERMANOVA *p* = 0.001). To capture more detail, we compared the prevalence and abundance differences of each enzyme subfamily in the metatranscriptomic samples. Nine subfamilies had higher prevalence (Fisher’s exact test *p* < 0.05, varying from 9% ∼ 17%) in the non-IBD group than the IBD group (**Supplementary Table 4**). We further identified 8 subfamilies (6 CYS, 1 CFAT, and 1 SDR) as significantly different in abundance (expression) profiles between the healthy controls and IBD cases (**Fig. 1f**, two-sided Mann-Whitney *U* test, adjusted *p* < 0.05). Notably, 7 of the 8 subfamilies had a higher prevalence (**Fig. 1f**, upper panel, Fisher’s exact test *p* < 0.05) and a higher abundance (**Fig. 1f**, lower panel, one-sided Mann-Whitney *U* test, adjusted *p* < 0.05) in the non-IBD group than in the IBD group.

We finally looked for metabolomic evidence in the differences of detectable SoLs, the products of the biosynthetic enzymes mentioned above, among IBD and non-IBD groups from publicly accessible metabolomics datasets. We expected that the increased expression of SoL biosynthetic enzymes would correspond with increased production of SoLs in non-IBD groups compared to IBD groups. Using metabolomics data from IBDMDB first, three metabolite features were assigned to sulfobacins B (SoL B), C (SoL C), and F (SoL F) by exact mass comparison with mass error less than 5 ppm (**Supplementary Data 3**). Indeed, we found that SoLs B, C, and F had significantly higher abundance in non-IBD groups than IBD groups (**Fig. 1g**, Wilcoxon rank sum test, *p* < 0.001). Furthermore, higher SoLs B, C, and F abundance was observed in non-IBD samples compared to both IBD subtypes: Ulcerative colitis (UC) and Crohn’s disease (CD) (**Extended Data Fig. 2a**, Wilcoxon rank sum test, *p* < 0.001). In another independent metabolomics dataset^37^, we found that the abundance of SoL B was also lower in IBD groups, consistent with the result generated from the first dataset (**Extended Data Fig. 2b**, Wilcoxon rank sum test, *p* < 0.001).

Thus, our metagenomic analysis reflected that SoL biosynthetic enzymes were more prevalent in the non-IBD group than the IBD group, metatranscriptomics suggested that these enzymes are more actively transcribed in the non-IBD group, and metabolomics indicated that representative SoLs are produced in higher abundance in the non-IBD group. Altogether, our findings establish a negative correlation directly between SoLs biosynthesis and IBD, consistent with the previously reported negative association between SoL-producers, namely *Alistipes* and *Odoribacter*, and IBD^17, 18^.

### A mouse study confirms the negative correlation between SoLs production and IBD progression

Encouraged by our informatically predicted negative correlation between SoL biosynthesis and IBD, we sought to experimentally validate our prediction using a mouse model of IBD. We used a well-established model of *Il10*-deficient (*Il10****^−/−^***) mice treated with the non-steroidal anti-inflammatory drug piroxicam, which has been shown to accelerate development of colitis through the disruption of the gut mucosal barrier^38, 39^. Stimulation of mucosal Toll-like receptors (TLRs) stemming from this mucosal barrier breakdown was another factor in our selection of this model, as we have previously shown that SoL A suppresses LPS-induced inflammation and LPS is well-known to activate TLR signaling^25, 40^. Accordingly, we successfully established the piroxicam/*Il10****^−/−^*** IBD model and observed that the colonic tissues were inflamed in the piroxicam-treated (IBD) group when compared to the control (pre-IBD) group as indicated by marked crypt hyperplasia, loss of goblet cells, submucosal edema, and immune cell infiltration into the lamina propria (**Fig. 2a,b, Supplementary Table 5**). Gross pathology analysis included qualitative evaluations of cecal atrophy, thickening of cecal and colon tissues, extent of content loss in the cecum, and diarrhea supported induction of IBD with piroxicam treatment (**Fig. 2c, Supplementary Table 6**).

**Fig. 2.**
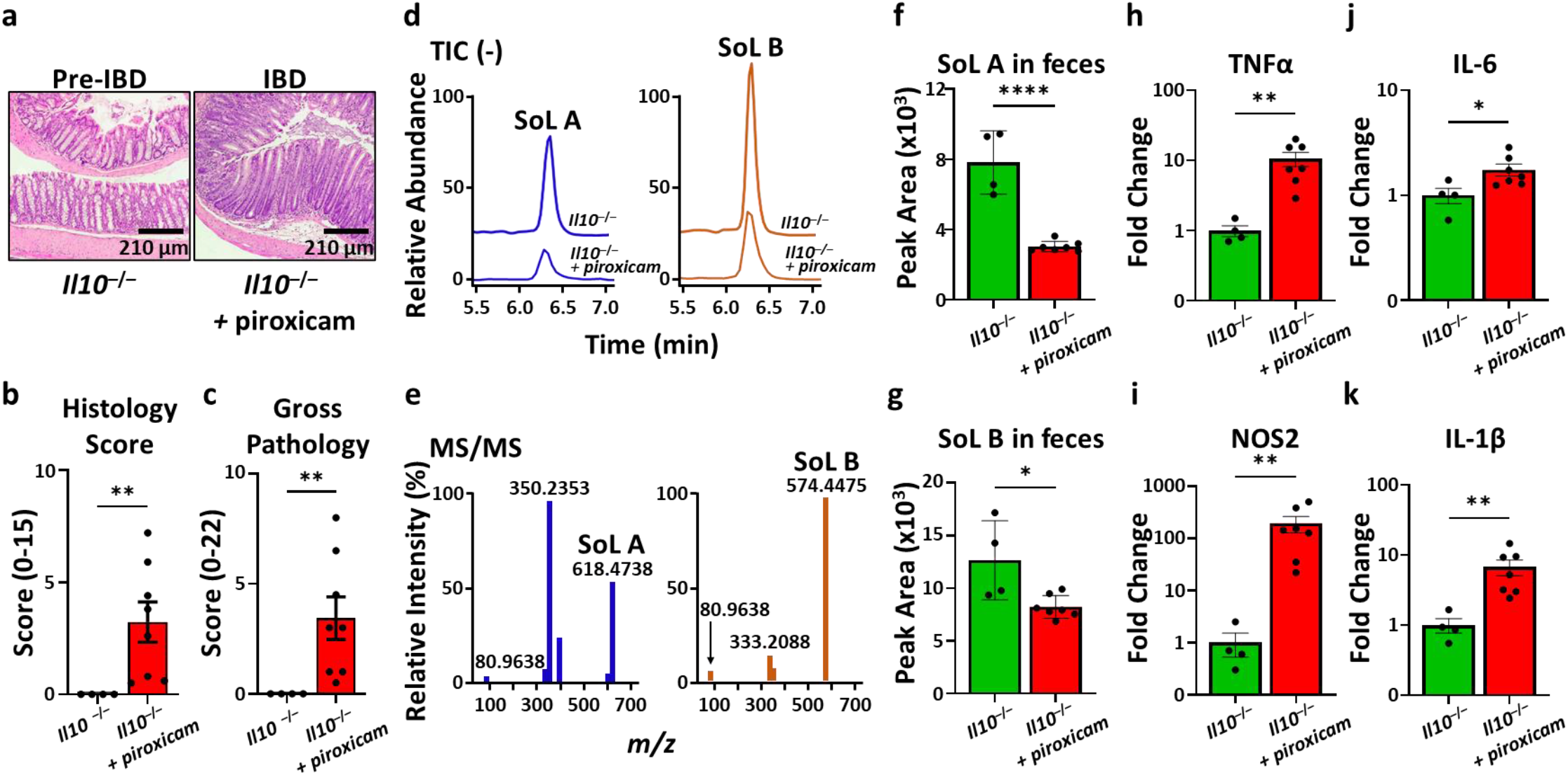
SoLs are decreased in a mouse model of IBD concurrent with increased concentration of inflammatory markers. **a**, Histological analysis of the mouse distal colon reveals that piroxicam treatment induced intestinal inflammation. **b** and **c**, Histology and gross pathology scores indicate induction of IBD in *Il10^−/−^* mice treated with piroxicam (*n* = 7, female) compared to pre-IBD control *Il10^−/−^* mice (*n* = 4, female), confirming the successful establishment of IBD model. The trends were consistent in male mice in another independent experiment using the same IBD model (**Extended Data Fig. 3**). **d**, Total ion chromatograms (TICs) obtained from HPLC-HRMS analysis of fecal pellet extracts from *Il10^−/−^* and *Il10^−/−^* + piroxicam mice reveal the presence of SoL A and SoL B. SoL abundances appear to be decreased in *Il10^−/−^* + piroxicam mice fecal pellets. **e**, MS/MS spectra of SoLs A and B confirm their identities based on the presence of the 80 *m/z* fragment characteristic of sulfonate-containing compounds and compared to literature fragmentation patterns^21, 25^. **f** and **g**, Peak areas were calculated using TICs obtained after MS/MS fragmentation and used to measure the abundance of SoLs A and B. Both SoLs A and B were significantly decreased in mice IBD samples. Significance was determined using Student’s *t*-test. **h**–**k**, Expression of inflammatory markers TNFα, NOS2, IL-6, and IL-1β were significantly increased in *Il10^−/−^* + piroxicam mice. Significance was determined using Mann-Whitney *U* test. Bars represent mean ± standard error. For all *p* values: ** 0.001 < *p* < 0 .01 and **** *p* < 0.0001.

To explore the link between SoLs production and IBD, we collected fecal material from IBD *Il10****^−/−^*** + piroxicam (*n* = 7, female) and pre-IBD control *Il10****^−/−^*** mice (*n* = 4, female), extracted metabolites, and measured the abundance of SoLs by targeted metabolomics using high performance liquid chromatography (HPLC)-high resolution mass spectrometry (HRMS) (**Fig. 2d**). We detected metabolites with *m/z* corresponding to major SoLs, specifically SoLs A and B, in all samples tested and unambiguously determined their identities by HPLC-MS/MS (**Fig. 2e**). We then determined that the abundances of both SoLs A and B were significantly decreased in piroxicam treated samples compared to *Il10****^−/−^*** control samples (**Fig. 2f,g**). This result confirms our above-described informatic analysis and directly establishes a negative correlation between SoLs production and IBD progression in the mouse model. In addition, we also observed significantly increased expression of the NF-κB-regulated inflammatory markers TNFα, NOS2, IL-6, and IL-1β in the IBD mouse group (Mann-Whitney *U* test, *p* ≤ 0.005; **Fig. 2h**–**k**), further indicating a negative correlation between SoL production and these inflammatory markers. Given our previously observed anti-inflammatory activity of SoL A against LPS^25^, a natural ligand of TLR4, this negative correlation suggests a potential role of SoLs in regulating IBD that may involve suppressing TLR4-mediated NF-κB activation. To eliminate any differences caused by sex, we performed another independent study with male mice using the same IBD model and observed the same negative correlation between SoLs production and IBD progression (**Extended Data Fig. 3**).

### Constant identification of SoLs’ contribution to immunomodulatory activity

We next examined the production of SoLs and its contribution to immunomodulatory activity in different human gut commensals. Unlike *C. gleum* F93 DSM 16776, which we experimentally investigated for its functional metabolites in relation to inflammatory activity^25^, the prolific SoL-producers *Alistipes* and *Odoribacter* had not yet been thoroughly chemically investigated to identify the biologically active components associated with remediation of IBD. In addition, *Alistipes* and *Odoribacter* produce a mixture of other SoLs^21^ and are likely to produce a multitude of other functional metabolites, both of which may complicate the potential immunomodulatory activity of these genera’s metabolites with respect to their bioinformatically predicted negative association with IBD. Thus, we conducted bioactive molecular networking of three *Alistipes* and two *Odoribacter* strains (**Supplementary Table 7**) to identify the constant contributor(s) to biological activity. We fractionated crude extracts of the *Alistipes* and *Odoribacter* strains and determined the biological activity of each fraction using a cell-based assay that measured the suppression of LPS-indued TNFα production (**Fig. 3a**). We simultaneously analyzed each fraction by untargeted HPLC-HRMS/MS to generate molecular networks using the Global Natural Products Social (GNPS) feature-based molecular networking (FBMN) pipeline^41^. We then correlated the relative expression of TNFα in each fraction with the relative peak area of molecular features across all fractions to generate a bioactivity score reflecting the contribution of specific features to the activity of the fractions. Bioactivity scores and relative peak areas were then mapped onto the molecular network to visualize these contributions. A representative bioactive molecular network generated from *Alistipes timonensis* DSM 27924 is presented in **Figure 3b**. The SoL-containing cluster contained the most abundant and most active molecular features, as indicated by the node size and color intensity compared to other clusters in the network. Additionally, this cluster contained several known SoLs but many more unannotated SoLs, suggesting that the family of biologically active SoLs is larger than what is currently known. In all other SoL-producers tested, we consistently identified SoLs as a major contributor in the active fractions of each strain (**Extended Data Fig. 4**). To exclude the possibility of observed SoL activity being influenced by LPS contamination, we confirmed the absence of leftover LPS in the SoL samples using a chromogenic LAL assay (**Supplementary Fig. 2**). Narrowing down the immunosuppressive activity of each of the strains to SoLs guided us to isolate pure SoLs A and B from *A. timonensis* DSM 27924 (structures confirmed by NMR spectroscopy; **Supplementary Tables 9 and 10**, **Supplementary Figures 3–14**), as well as from each of the other *Alistipes* and *Odoribacter* strains tested. We thus reinforced the contribution of this class of lipids to the observed biological activity of *Alistipes* and *Odoribacter*.

**Fig. 3.**
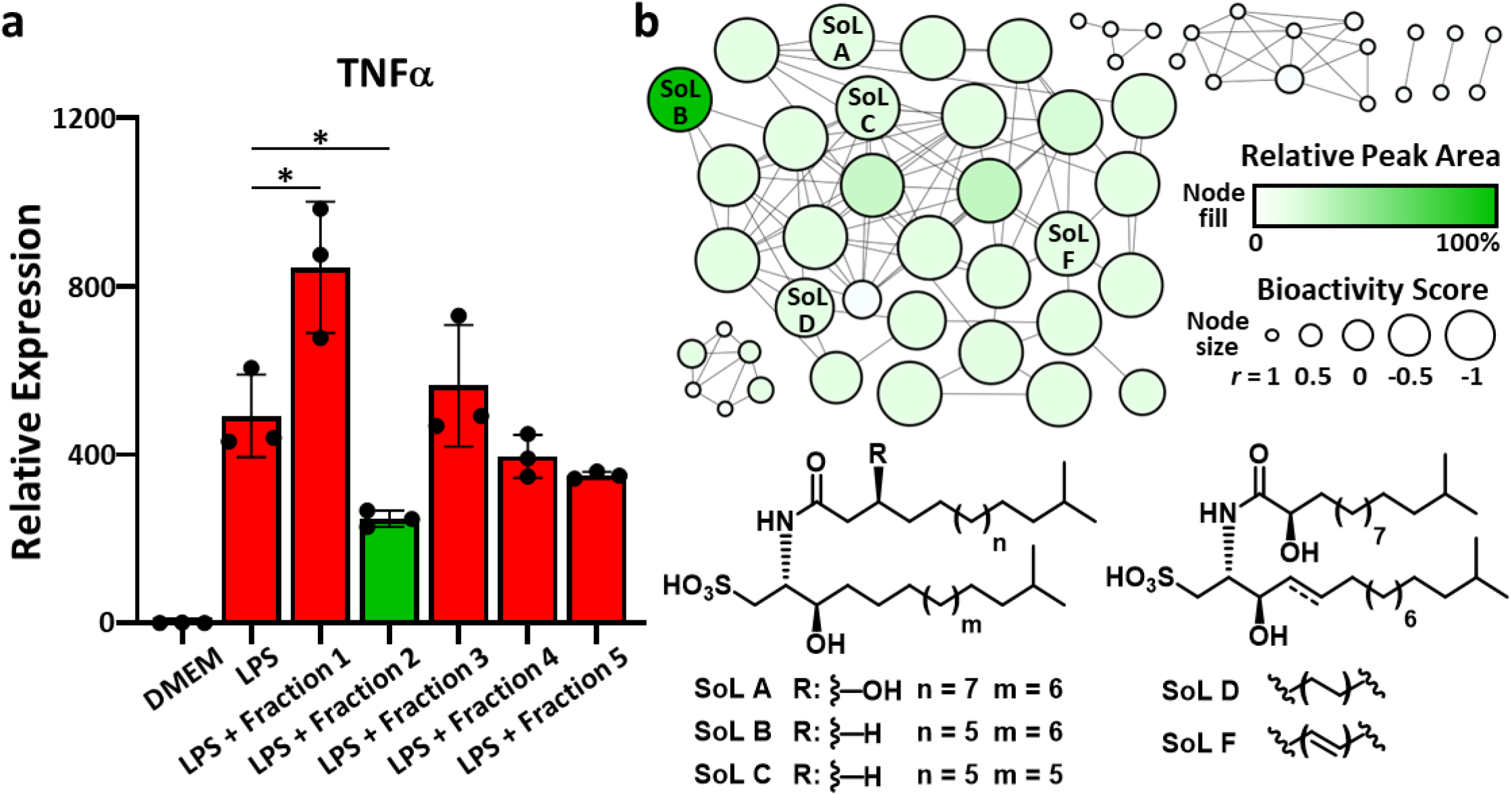
Bioactive molecular networking leads to the identification of SoLs as major bioactive components of a SoL-producer. **a**, A crude extract of *A. timonensis* DSM 27924 was separated based on polarity into 5 fractions. Each fraction was used in an *in vitro* cell-based assay to measure its respective capacity to suppress LPS-induced expression of TNFα. Fraction 2 was found to have the most significant anti-inflammatory effect compared to LPS. All fractions were compared to LPS for statistical significance with only fraction 1 and 2 showing significant change. Fractions 3, 4, and 5 showed no significant change. Statistical significance was determined using Student’s t-test. Bars represent mean ± standard error. For all *p* values: * 0.01 < *p* < 0.05. **b**, Untargeted HPLC-HRMS/MS was used to construct a molecular network for each fraction through GNPS FBMN. Relative peak area of each molecular feature in fraction 2 was mapped to the color of the nodes with more abundant features increasing from white to green. Bioactivity score was mapped to the node size with larger nodes indicating stronger negative correlations. Several known SoLs were annotated in this cluster and their structural variations are illustrated, further demonstrating that SoLs as a family of molecules contribute to the observed suppression of LPS-induced TNFα expression.

### SoL A mediates dual immunomodulatory activity through TLR signaling

While the causes of IBD remain largely unknown, IBD progression has been linked to aberrant TLR signaling^40^. TLRs are pattern recognition receptors (PRRs) which initiate a variety of host processes, especially inflammatory responses, through the recognition of pathogen-associated molecular patterns (PAMPs) and other non-pathogenic microbial factors^40, 42, 43^. Specifically, TLR2 and TLR4 are well-known to recognize PAMPs in the gut microbiome^40^. In addition, their expression is significantly increased in IBD pathogenesis, reflecting a state of aberrant activation^40, 43^. Thus, we expected that SoLs may interact with TLR4 or TLR2 to mediate their immunomodulatory activity. We treated primary mouse macrophages with SoL A (as a representative of SoLs) either alone or together with LPS (an agonist of TLR4) or Pam3CSK4 (an agonist of TLR1/2) and measured the expression of three inflammatory cytokines (IL-6, TNFα, and IL-1β). By itself, SoL A exhibited a mild to moderate effect on the expression of pro-inflammatory cytokines compared to control (**Fig. 4**), generally consistent with our previous finding^25^. As expected, the TLR ligands, LPS and Pam3CSK4, both showed significant induction of all three cytokines compared to control (Student’s *t* test, *p* ≤ 0.0001) (**Fig. 4**). Notably, SoL A was found to significantly suppress the expression of all three cytokines induced by LPS (*p* ≤ 0.0001) (**Fig. 4**). Together, SoL A’s mild pro-inflammatory activity by itself and primarily strong inhibition against LPS-induced inflammation constitute its dual immunomodulatory activity. SoL A also inhibited Pam3CSK4-induced IL-6 and TNFα to a smaller extent while increasing IL-1β expression induced by Pam3CSK4 (*p* ≤ 0.05) (**Fig. 4**). This result indicates that SoL A primarily affects LPS-induced inflammation and implies that interaction with TLR4 may be involved in SoL A’s mechanism of action. Interestingly, SoL A’s partial suppression of Pam3CSK4-induced inflammation suggests that SoL A-related anti-inflammatory activity may also extend to the TLR1/2 pathway, albeit to a lesser extent, and warrants further investigation.

**Fig. 4:**
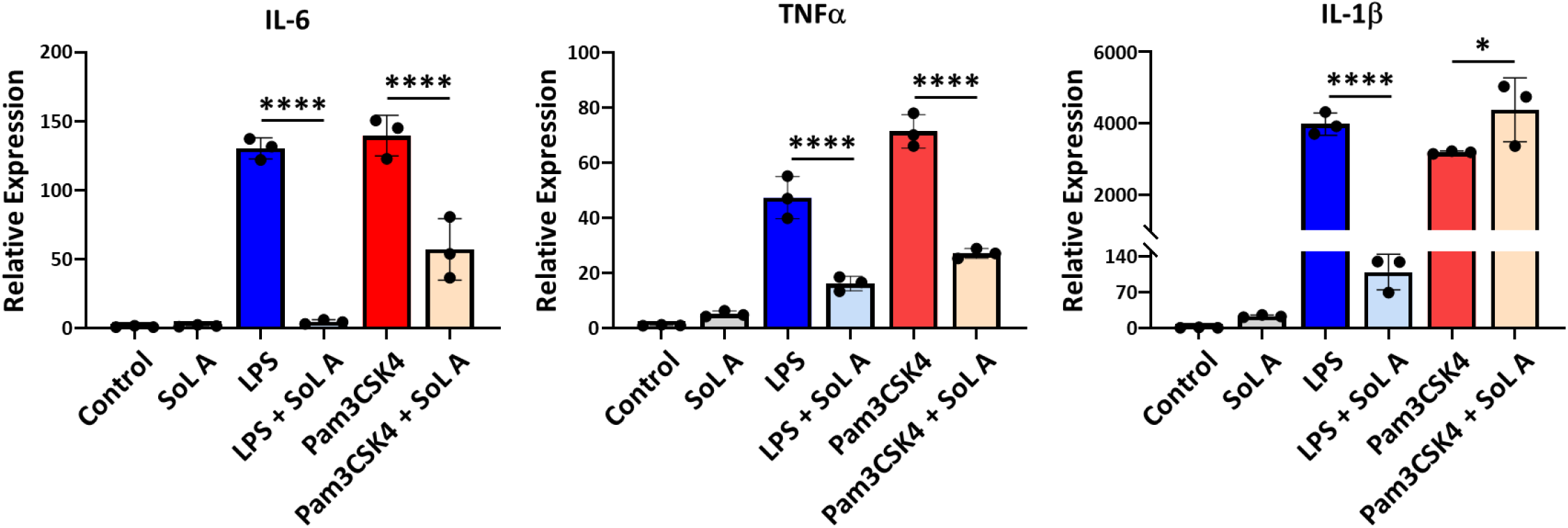
SoL A primarily suppresses LPS-induced TLR4 activation. Mouse peritoneal macrophages were treated with SoL A (10 μM), LPS (100 ng/mL), and Pam3CSK4 (500 ng/mL), either alone or in combination for 6 hours. RT-qPCR analysis revealed that SoL A induces a mild pro-inflammatory effect compared to control but significantly suppresses LPS-induced cytokine expression levels and only partially suppresses Pam3CSK4-induced cytokine expression. Bars represent mean ± standard error. For each treatment, *n* = 3. Significance was determined using Student’s *t* test: * *p* < 0.05, **** *p* < 0.0001.

### Molecular docking and ELISA displacement assay suggest SoLs binding to TLR4/MD-2 complex

LPS stimulation of TLR4 occurs through a series of interactions ultimately resulting in LPS binding to MD-2, which forms a complex with TLR4 and induces dimerization to initiate signaling^44–46^. The TLR4/MD-2 heterodimer recognizes structurally diverse LPS molecules, giving it flexibility to detect different LPS-related PAMPs in the human gut microbiome^46^. Interestingly, the TLR4/MD-2 complex was recently found to recognize human sulfatides, sphingolipid derivatives which bear a sulfated saccharide head group and dual acyl chains, presumably mimicking the disaccharide core and multiple acyl chains of LPS^47^. Comparing the chemical structure of SoL A to those of sulfatides and lipid A (the immunogenic portion of LPS) (**Fig. 5a**), we noted structural similarity in the negatively charged head groups and multiple acyl chains. We thus considered if multiple molecules of SoL A might bind to MD-2 in a similar configuration as sulfatides and lipid A. Inspired by sulfatides that bind in triplicate to MD-2^47^, we used molecular docking to model the binding of three molecules of SoL A to MD-2. Our analysis predicted three molecules of SoL A indeed bind in the hydrophobic pocket of MD-2 (**Fig. 5b**), where lipid A is known to bind, with a docking score of -8.9 kcal/mol, better than that of lipid A which had a docking score of -6.2 kcal/mol. Additionally, SoL A was predicted to make hydrophobic contacts with several amino acids including I117, F119, I52, and F121 (**Supplementary Table 8**), all of which are also reported to contact the acyl chains of lipid A^46^. Notably, SoL A is also predicted to contact residues including R264 and R90 (**Supplementary Table 8**), consistent with contacts between these residues and the phosphate groups of lipid A^46^. This suggests that SoL A may bind directly to the TLR4/MD-2 complex and possibly compete for binding with LPS, allowing it to suppress LPS-induced activation of the TLR4 pathway. A critical aspect of lipid A binding to MD-2 is the exclusion of one acyl chain from the hydrophobic pocket of MD-2 which forms a bridge with TLR4 and is involved in inducing dimerization^46^. Likewise, we observed one acyl chain of SoL A excluded from the hydrophobic pocket in our docking analysis (**Fig. 5b**), further suggesting that SoL A mimics LPS as a ligand for TLR4. After successfully docking of SoL A, we tested SoL B which lacks an extra hydroxy group (**Fig. 5a**) that may increase its interactions with the hydrophobic binding pocket of MD-2. Our docking analysis indeed showed that SoL B also binds to MD-2 (**Fig. 5b**), with similar contacts as SoL A (**Supplementary Table 8**) but higher affinity (docking score of -9.6 kcal/mol) as predicted.

**Fig. 5:**
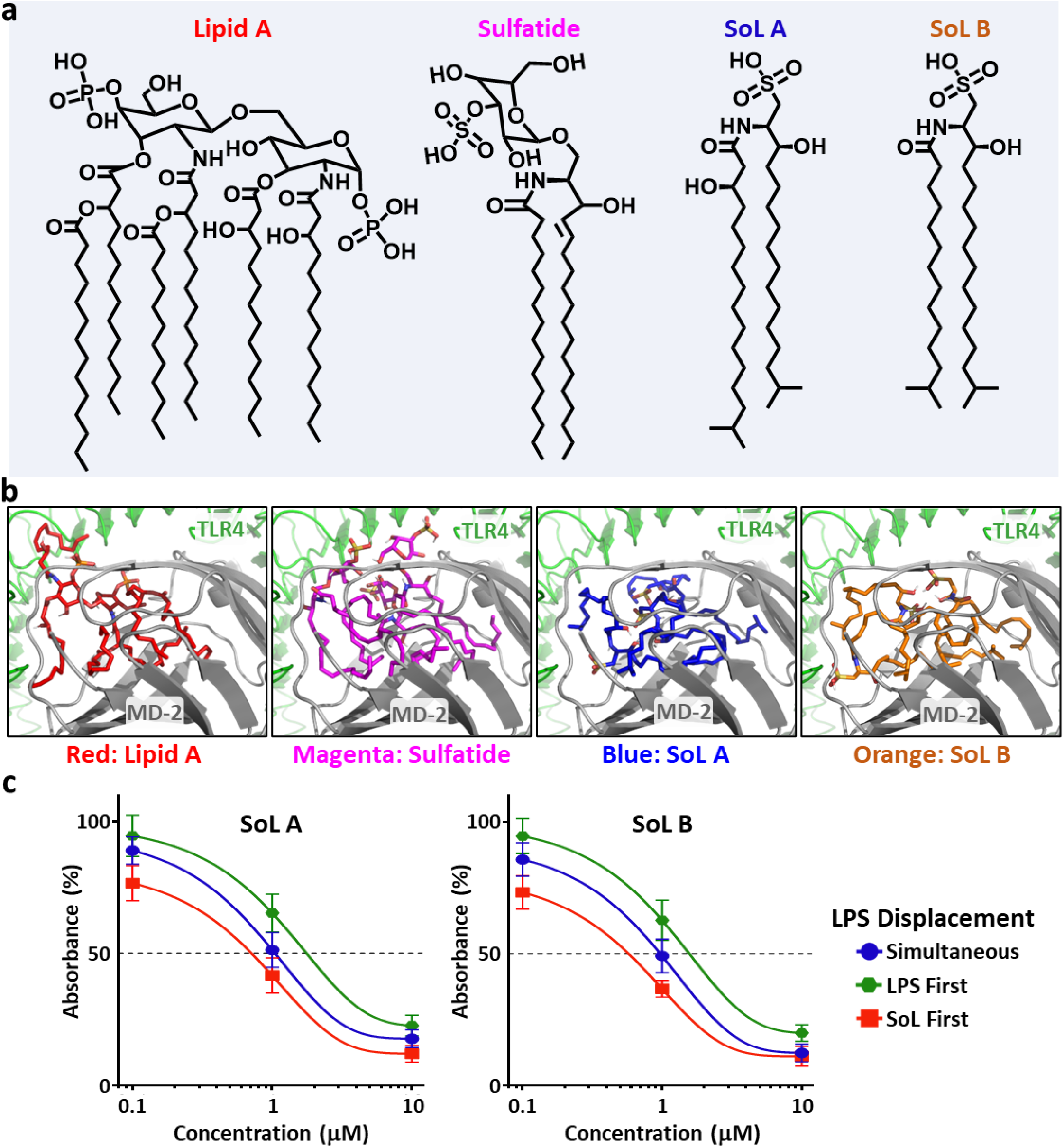
SoLs bear structural similarity to both lipid A and sulfatide and bind with MD-2 to block LPS binding. **a**, Chemical structures of immunogenic lipid A, derived from LPS, sulfatide, SoL A, and SoL B illustrating structural similarity in multiple acyl chains and negatively charged head groups. **b**, Molecular docking of lipid A (red), sulfatide (magenta), SoL A (blue), and SoL B (orange) into the hydrophobic pocket of MD-2 in complex with TLR4. Three molecules of SoLs A and B were used in molecular docking experiments to mimic the six acyl chains of lipid A as inspired by sulfatides^47^. **c**, ELISA displacement assay used to measure the binding behavior of SoLs A and B in competition with 1 ng/mL LPS, a natural ligand of the TLR4/MD-2 complex. Compounds were added either simultaneously (blue), LPS first (green), or SoL first (red).

To experimentally determine if SoLs A and B bind to MD-2 and to what extent the SoLs compete with LPS for binding to MD-2, we conducted an ELISA-based displacement assay. Taking advantage of biotinylated LPS, which retains the activity of unconjugated LPS^48^, we measured absorbance generated by an HRP-linked streptavidin probe to measure the relative amount of MD-2 which was bound with biotin-LPS as opposed to MD-2 bound with SoL A or B. We administered 0.1, 1.0, and 10 μM concentrations of SoL A or B and 1 ng/mL biotin-LPS to MD-2 in three sequences: 1) SoL first followed by LPS 1 hour later, 2) LPS first followed by SoL 1 hour later, and 3) both SoL A or B and LPS at the same time. After 1 hour of incubation, we found that at all concentrations, when SoL A or B was added first, there was a marked decrease in percent absorbance as compared to when LPS was added first and when the two compounds were added together (**Fig. 5c**). This suggests that SoLs A and B both bind and occupy some sites of MD-2, preventing LPS from fully binding when it is added 1 hour after SoL A or B. Furthermore, when moving from low to high concentrations of SoL A or B, we observed that the percent absorbance decreased dramatically. This indicates that with increasing concentration of SoL A or B, less LPS binds to MD-2, implying that SoLs indeed compete with LPS for binding to MD-2. Taken together, these results indicate that SoLs A and B can bind directly to MD-2 and more importantly compete with LPS for binding to this target, thus providing a potential molecular mechanism underlying SoL A’s pro-inflammatory activity by itself as well as its strong activity in suppressing LPS-induced inflammation which likely also expands to other members of SoLs family.

### SoLs suppress LPS-induced TLR4 signaling to regulate macrophage polarization

Upon LPS binding, TLR4 initiates downstream signaling, such as through the NF-kB and MAPK pathways, resulting in the induction of inflammatory cytokine expression^43^. If a SoL binds to MD-2 to suppress LPS-induced inflammation, this would block activation of the TLR4 pathway. Therefore, we investigated whether addition of SoL A or B affected the phosphorylation of TLR4-downsteam signaling molecules, ERK1/2 and p38, and the degradation of IκBα, which are critical for LPS-induced cytokine expression^43^ (**Fig. 6a**). We treated macrophages with LPS in the presence of increasing concentrations of SoL A or B (from 0 to 20 μM), then performed western blot analysis to examine the TLR4-downstream signaling pathways. We found that both SoLs reduced LPS-induced phosphorylation of p38 and ERK1/2 in a concentration-dependent manner. At the concentration of 20 μM, SoLs A and B almost completely blocked LPS-induced phosphorylation of p38 and ERK1/2 (**Fig. 6b**). Western blot also showed that SoLs concentration-dependently suppressed LPS-induced IκBα degradation (**Fig. 6b**). These results support that SoLs exert their anti-inflammatory effect by blocking LPS-mediated phosphorylation of downstream TLR4 proteins, effectively negating LPS activation of the TLR4 pathway. Also, SoL A or B alone at the concentration of 20 μM slightly enhanced the phosphorylation of certain signaling molecules (e.g., ERK1/2) compared to control, consistent with our observations that SoLs alone induced a mild pro-inflammatory effect on cytokine expression (**Fig. 4**) and supporting the dual immunomodulatory activity of SoLs which provides further opportunities to regulate homeostatic immune responses.

**Fig. 6:**
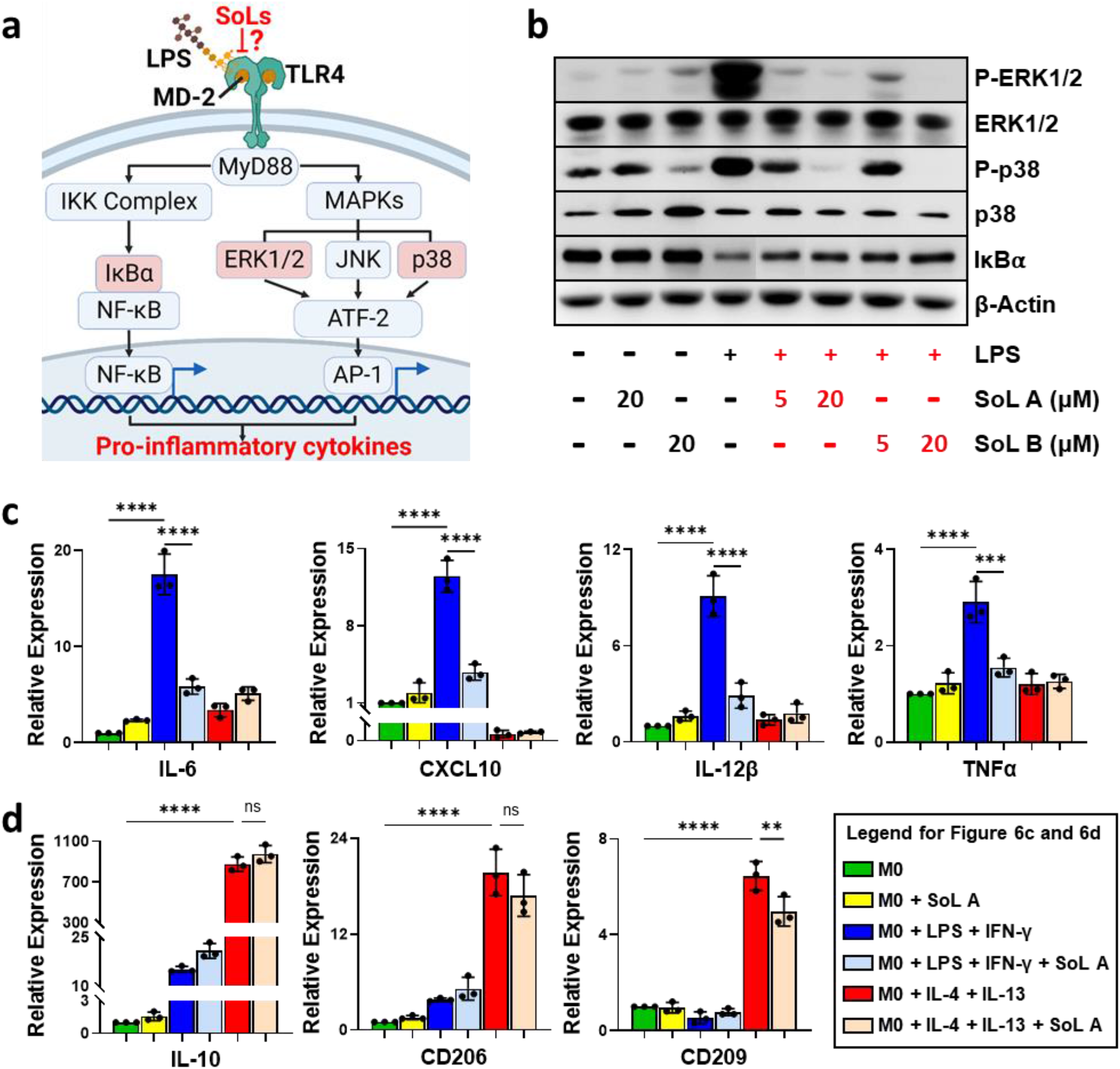
SoLs suppress LPS-induced activation of TLR4 signaling pathway and macrophage M1 polarization. **a**, Simplified pathway of TLR4 activation by LPS, highlighting proposed inhibition by SoLs competing for LPS binding. Proteins IκBα, ERK1/2, and p38 downstream of TLR4 which were selected for analysis are highlighted in red rectangles. **b**, Western blot analysis of protein levels of IκBα as well as total and phosphorylated ERK1/2 and p38, after treatment with LPS (100 ng/mL) with or without various concentrations of SoL A or B. The housekeeping gene β-Actin was used as a loading control. **c** and **d**, THP-1-derived macrophages were treated with LPS + IFN-γ or IL-4 + IL-13 to polarize to M1 or M2 macrophages, respectively. Relative expression of markers IL-6, CXCL10, IL-12β, and TNFα (compared to M0) indicate that SoL A (10 μM) had a significant effect on suppressing M1 polarization (**c**); and relative expression of markers IL-10, CD206, and CD209 (compared to M0) indicate that SoL A had no significant effect on M2 polarization (**d**). For all treatments, *n* = 3 and significance was determined using one-way ANOVA: ** 0.001 < *p* < 0.01, *** 0.0001 < *p* < 0.001, and **** *p* < 0.0001.

Because TLR4 signaling leads to macrophage polarization which has been shown to contribute to IBD^49–51^, we also examined the effects of SoL A on macrophage polarization. We treated THP-1 monocytes with IFN-γ and LPS to induce M1 polarization or IL-4 and IL-13 to induce M2 polarization. Successful induction of M1 and M2 polarization was confirmed by morphology changes and subsequent RT-qPCR quantification of cytokine profiles. When 10 μM of SoL A was added alongside the respective inducing agents, our relative cytokine expression results showed that SoL A significantly reduced the production of M1-polarized macrophage markers IL-6, CXCL10, IL-12β, and TNFα, compared to macrophages treated without SoL A (**Fig. 6c**) but had a mostly non-significant effect on M2 polarization (**Fig. 6d**). This suggested that SoL A suppresses macrophage M1 polarization, which supports our aforementioned result that SoLs interfere with TLR4 signaling potentially leading to inhibition of TLR4-mediated IBD.

## Discussion

Taking advantage of our unique biosynthetic enzyme-guided disease correlation approach, the results of this study have described two critical points, summarized here and discussed in further detail below. First, we have directly connected the biosynthesis and production of a class of abundant yet underexplored human microbial metabolites, SoLs, to IBD, an existing human health condition with complex and poorly understood etiology, followed by a mouse model of IBD to validate this informatically predicted negative correlation. Second, we have revealed that SoLs A and B, two representative gut microbial SoLs, modulate host immune responses through the TLR4/MD-2 complex and inhibit LPS-induced TLR4 activation, which provides a mechanistic explanation for SoLs’ potential protective activity against IBD.

Through both bioinformatic and cheminformatic analyses, we revealed that the expression of SoL biosynthetic enzymes and abundance of SoLs in the gut metabolome are negatively correlated with IBD incidence in humans. Our IBD model with piroxicam treated *Il10****^−/−^*** mice supported this negative correlation with a concurrent negative association between SoL production and TLR-4-related inflammatory markers TNFα, NOS2, IL-6, and IL-1β. Our findings were consistent with literature reports which have shown that the bacterial genera of abundant SoLs production, *Alistipes* and *Odoribacter*, are associated with the remediation of IBD symptoms^19, 20^. Considering that these SoL-producers are commensal members of the human gut microbiome^21, 52^, their constant production of SoLs in the human gut may help to maintain intestinal immune homeostasis, thereby preventing IBD. Both *Alistipes* and *Odoribacter* are also known to produce short chain fatty acids (SCFAs), which have been implicated in reducing intestinal inflammation^14, 20, 52–54^. Besides SCFAs, there are other gut microbial functional metabolites such as secondary bile acids and indole derivatives which have also been linked to modulating host inflammation and immunity^55–59^. Thus, the role SoLs play individually and/or synergistically with other factors in IBD pathogenesis remains interesting and awaits further investigation.

Towards understanding the mechanism of the immunomodulatory activity of SoLs, we showed that the representative SoLs A and B primarily target the TLR4 pathway and that they both block LPS binding to MD-2 to suppress TLR4 signaling. Our analysis indicated that SoL A preferentially suppresses LPS-induced inflammation as compared to Pam3CSK4-induced inflammation, suggesting that TLR4 activation is more strongly affected by SoL A than TLR1/2 activation. The selectivity for TLR4 may stem from SoL A’s structural similarity to LPS, a hypothesis supported by the recently reported human TLR4 ligands, sulfatides, which share highly similar structural features to SoLs^47^. As microbial functional metabolites, it is reasonable that SoLs are directly recognized by TLR4, which has evolved specifically to recognize PAMPs^60^. Our molecular docking also suggested SoLs A and B’s recognition by TLR4, indicating that three molecules of either SoL indeed bind with stronger predicted binding score compared to lipid A and make contacts with important amino acids in the pocket of MD-2, consistent with lipid A binding as well as structurally-related sulfatides^46, 47^. Our ELISA-based displacement analysis then confirmed that both SoLs A and B directly bind to MD-2, block LPS binding when added prior to LPS, and could displace bound LPS from MD-2 at higher concentrations. We further demonstrated that increasing concentration of SoLs A and B suppressed LPS-induced TLR4 signaling pathways in a dose-dependent manner and SoL A significantly suppressed M1 polarization of macrophages, further indicating their capacity to reduce downstream TLR4 signaling responses. Notably, increased numbers of M1 macrophages is a characteristic feature of IBD^51^ and this suppressive effect may represent one explanation how SoL-producing bacteria are able to remediate symptoms of IBD. Taken together, our results represent the first report of SoLs A and B’s binding to MD-2 and establish that SoLs likely mediate their dual immunomodulatory activity by occupying the hydrophobic binding pocket of MD-2 (pro-inflammatory by SoLs alone) but primarily blocking LPS binding to the TLR4/MD-2 complex (anti-inflammatory against LPS-induced inflammation). This discovery suggests SoLs’ mechanistic role in regulating a multitude of TLR4-related inflammatory conditions, most notably IBD which is associated with dysregulation of TLRs, especially TLR4^26–29^, leading to aberrant macrophage activation^51, 61^.

While many studies have focused on the association of certain microbial strains with specific diseases by analyzing the abundance and distribution of strains^17, 18, 62–65^, our biosynthetic enzyme-guided disease correlation approach has shown that the presence and expression of biosynthetic enzymes corresponding to functional metabolites can be directly correlated with human health conditions, effectively shifting the microbe-focused perspective to a functional metabolite-based molecular perspective. By applying this approach, we have identified a negative relationship between SoL biosynthesis and IBD directly from patients’ data followed by verification using an IBD mouse model, which has guided us to further reveal a molecular target and a potential mechanistic explanation for the protective effect that SoLs and SoL-producing bacteria exert against IBD. We are now further characterizing the effect of SoL biosynthesis on IBD pathogenesis through in-depth *in vivo* studies using purified SoLs as well as developing isogenic mutants of SoL-producing bacteria for mouse colonization studies. With the exponentially increasing availability of human disease omics data, we expect our approach described in this study to be widely applicable to uncovering the molecular mechanisms of other intricate host-microbe interactions.

## Methods

### Identification of SoL biosynthetic enzymes from human gut bacterial reference genomes

We collected experimentally validated enzymes involved in SoL biosynthesis as reference amino acid sequences of CYS, CFAT, and SDR (**Supplementary Table 1**). Reference amino acid sequences were used as seed sequences to search for homologs in the human gut bacterial reference genomes using the DIAMOND blastp model^66^ with an e-value threshold of 10^-5^. We then investigated the taxonomic distribution of the resulting homolog sets based on the taxonomy annotation for each genome^30^. To prioritize SoL biosynthetic enzymes, the homologs of CFATs and CYSs from genomes which encode copies of CFAT, CYS and SDR were first used to generate sequence similarity networks with experimentally validated CFAT and CYS at a threshold of 50% similarity using MMseqs2^67^. The prioritized CFATs and CYSs meet co-occurrence with SDRs in the same genome. The filtered enzymes were further subjected to Pfam domain analysis by hmmsearch (HMMER v3.3) with default parameters against the Pfam-A database (v33.1). Enzymes containing corresponding Pfam domains with hit score > 50 were selected as prioritized SoL biosynthetic enzymes. Prioritized enzymes were used to generate CYS, CFAT, and SDR subfamilies using sequence similarity networks with at least 90% similarity by MMseqs2 clustering^67^. Maximum-likelihood trees were generated using the representative genome of each species by GTDB-Tk (v2)^68^. Finally, we selected a representative genome for each species to display their SoL biosynthetic potential and annotate their phylogenetic trends using iTOL^69^.

### Quantification of enzyme abundances in metagenomic and metatranscriptomic samples

Metagenomic and metatranscriptomic whole-genome sequencing datasets of human gut microbiomes related to IBD were downloaded from the NCBI Sequence Read Archive (SRA) (SRA Accessions: PRJNA398089 and PRJNA389280). For both metagenomic and metatranscriptomic samples, reads were quality filtered and adapter removed using *bbduk.sh* with the following parameters: qtrim=rl ktrim=r mink=11 trimq=10 minlen=40 (read quality cutoff is 10, read length cutoff is 40). High-quality reads were mapped to the nucleotide sequences of corresponding contigs containing the SoL biosynthetic genes using BWA mem algorithm^70^ with default parameters. The reads mapped to genes were counted by featurecounts^71^ with the following parameters: -f -t CDS -M -O -g transcript_id -F GTF -s 0 -p --fracOverlap 0.25 -Q 10 –primary. Enzymes encoded or expressed in at least 5% of samples were considered as common distribution in humans and were included in the comparative analysis. Transcripts per million (TPM) were calculated for each SoL biosynthetic enzyme gene. The abundance of each cluster was calculated by the sum of the relative abundances of all genes in the cluster. Beta diversity was performed to quantify the prevalence and relative abundance differences in the overall composition of SoL biosynthetic enzymes between the IBD and the control groups. PERMANOVA was performed to show the encoding and expression profile differences of SoL biosynthetic enzymes between IBD and control groups. Both beta diversity and PERMANOVA were performed using the R package vegan^72, 73^. To explore the differences between IBD and control groups of single SoL biosynthetic enzyme subfamilies, we used the Shapiro–Wilk test to evaluate the normality of a specific gene cluster’s relative abundance. We then calculated the significance of relative abundance between healthy and IBD individuals using either two-sample Student’s t-test (for normally distributed data) or two-sample Wilcoxon rank sum test (for not normally distributed data). Significance tests were performed in Python using packages Pandas and SciPy^74–76^. The Benjamini-Hochberg method was used to adjust p-values to correct for multiple testing^77^. SoL biosynthetic enzyme subfamilies were considered differential if the adjusted p-value was less than 0.05. For differential Sol biosynthetic enzymes, we performed a two-sided Fisher’s exact test to explore their difference in prevalence across IBD and non-IBD groups.

### Analysis of publicly available metabolomics datasets

Two publicly available metabolomics datasets were downloaded from IBDMDB (https://ibdmdb.org/tunnel/public/summary.html) and Metabolomics Workbench (http://www.metabolomicsworkbench.org). Processed metabolomic feature tables were used to search for SoL related metabolites (SoLs A–F). Search parameters were set to the exact mass of SoLs A–F using a 5 ppm match tolerance for parent ions under negative mode. The absolute intensity of relative abundance from resulting matches were used to calculate differential abundances between non-IBD and IBD samples. The Wilcoxon rank sum test was used to measure statistical significance.

### Animals

*Il10^−/−^* mice were originally purchased from The Jackson Laboratory. Mice were maintained in specific pathogen free conditions at the University of South Carolina and were maintained on a 12-hour light/dark cycle with unlimited access to water and food (Envigo 8904). All animal protocols were approved by the University of South Carolina Institutional Animal Care and Use Committee.

### Piroxicam-accelerated *Il10^−/−^* mouse IBD model and analysis

At 8-12 weeks of age, male and female *Il10^−/−^* mice (*n* = 7 per sex) were switched to a diet containing 100 ppm of piroxicam (Sigma) (Envigo, TD.210442) to induce colitis development as previously described^39^. Male (*n* = 3) and female (*n* = 4) *Il10^−/−^* mice were maintained on the control diet for the duration of the experiment to serve as pre-IBD controls. Mice were euthanized at 18 days to collect tissues for assessment and intestinal contents for quantification of SoLs. At necropsy, IBD severity was first grossly assessed, which included qualitative evaluations of cecal atrophy (0–5), thickening of cecal (0–5) and colon tissues (0–5), extent of content loss in the cecum (0–4) and diarrhea (0–3). For histopathology, segments of the colon were first washed in PBS and then fixed in 10% neutral buffered formalin. The tissues were embedded in paraffin, cut into 5-mm sections, and stained with hematoxylin and eosin (H&E) at the Instrumentation Resource Facility at the University of South Carolina School of Medicine. Inflammation scores of colon sections were blindly assessed as previously described^78^ using an Echo Revolve light microscope and accompanying software. Briefly, IBD severity was assessed based on the following histopathological features: length measurements in microns of crypt hyperplasia converted to a score from 0-4, qualitative assessment of goblet cell loss (0–5), crypt abscesses per 10X field counts converted to a score from 0-4, and qualitative assessment of submucosal edema (0-3). RNA isolations and RT-qPCR were performed as previously described^79^. Briefly, RNA was isolated from snap frozen colon tissues using the TriZol method (Thermo Fisher Scientific). cDNA was synthesized using SuperScript III reverse transcriptase (ThermoFisher Scientific). qRT-PCR was performed at the Functional Genomics Core at the University of South Carolina. The relative abundance of mammalian mRNA transcripts was calculated using the delta delta CT method and normalized to *Eef2* levels. The oligonucleotides used for qRT-PCR were: *Eef2* forward: TGTCAGTCATCGCCCATGTG, reverse: CATCCTTGCGAGTGTCAGTGA; *Tnfa* forward: AGCCAGGAGGGAGAACAGAAAC, reverse: CCAGTGAGTGAAAGGGACAGAACC; *Nos2* forward TTGGGTCTTGTTCACTCCACGG, reverse: CCTCTTTCAGGTCACTTTGGTAGG. Fecal samples were collected and immediately flash frozen and stored at -80 C. For SoL extraction and quantification, frozen fecal samples were lyophilized to remove remaining water and subsequently resuspended in methanol (ThermoFisher Scientific). Fecal sample suspensions were vortexed for 1 minute prior to sonication for 10 minutes. The methanol extract was collected by centrifugation at 20,000 x *g* for 10 minutes and dried under a gentle stream of nitrogen. The resulting residue was then redissolved in methanol + 0.1% ammonium hydroxide (ThermoFisher Scientific) and filtered through a 0.22 μm filter prior to analysis. High-resolution mass spectra were collected using a ThermoFisher Scientific Q-Exactive HF-X hybrid Quadrupole-Orbitrap mass spectrometer using electrospray ionization in negative mode. Liquid chromatography used a ThermoFisher Scientific Vanquish HPLC coupled to the aforementioned mass spectrometer. LC was performed using a Waters Xbridge BEH C18 XP column (2.1 x 100 mm) with mobile phases A (water + 0.1% ammonium hydroxide) and B (acetonitrile + 0.1% ammonium hydroxide) in a gradient starting from 10% B and increasing to 100% B over 5 minutes, hold at 100% B for 2.5 minutes, then re-equilibration at 10% B for 2.5 minutes. MS scans were obtained in the orbitrap analyzer which was scanned from 500 to 2000 m/z at a resolution of 60,000 (at 200 m/z). MS data was analyzed by Thermo Xcalibur (4.2.47).

### Anaerobic culture and bioactive molecular networking

Three *Alistipes* and two *Odoribacter* strains (**Supplementary Table 7**) were cultured in Reinforced Clostridial Medium (RCM, BD Biosciences) under anaerobic conditions at 37°C. After three days of growth, cultures were harvested by centrifugation at 12,000 x *g* for 30 minutes. The resulting cell-free supernatant was extracted with an equal volume of methyl ethyl ketone and the cell pellets were extracted by resuspension in methanol and sonication before both extracts were combined and concentrated *in vacuo*. The combined crude extract was then fractionated on a silica gel column using a stepwise gradient of dichloromethane and methanol (DCM:MeOH; 15:1, 7:1, 5:1, 3:1, 1:1). Each fraction was then used in an *in vitro* cell-based assay measuring the suppression of LPS-induced TNFα expression. Simultaneously, samples of the fractions were subjected to untargeted HPLC-HRMS/MS as described above. MS/MS was conducted using data-dependent acquisition with a resolution of 30,000, isolation window of 2.0 *m/z*, and dynamic exclusion time of 15 seconds. HPLC-HRMS/MS data was processed using MZmine3 following the GNPS FBMN workflow with minimal changes^80^. Molecular networks were constructed using the quickstart GNPS FBMN setting with no changes^41^. Bioactivity scores were assigned using a custom R script which calculated Pearson correlation coefficients between each molecular feature and the activity of each fraction^81^. Finally, bioactive molecular networks were visualized in Cytoscape v3.9.1^82^.

### Purification of SoLs A and B

Fractions containing SoLs were further purified by Sephadex LH-20 run in 1:1 DCM:MeOH. Finally, pure SoLs A and B were isolated by semi-preparative scale HPLC running an isocratic solvent composition of 47% water (with 0.1% ammonium hydroxide) and 53% acetonitrile (with 0.1% ammonium hydroxide) on a ThermoFisher Scientific Ultimate 3000 semi-preparative scale HPLC equipped with a Waters Xbridge Prep C18 5 μm OBD column (19 x 100 mm). ^1^H, ^13^C, ^1^H-^13^C HSQC, ^1^H-^13^C HMBC, and ^1^H-^1^H COSY NMR spectra for SoLs A and B were acquired in methanol-d4 on a Bruker Avance III HD 400 MHz spectrometer with a 5 mm BBO 1H/19F-BB-Z-Gradient prodigy cryoprobe. Data were collected and reported as follows: chemical shift, integration multiplicity (s, singlet; d, doublet; t, triplet; m, multiplet), coupling constant. Chemical shifts were reported using the methanol-d4 resonance as the internal standard for ^1^H-NMR methanol-d4: δ = 3.31 ppm and ^13^C-NMR methanol-d4: δ = 49.0 ppm. Pure SoLs A and B were confirmed to be free of LPS using a Chromogenic Endotoxin Quant Kit (Pierce).

### Preparation and treatment of macrophages

Primary mouse macrophages were prepared by first introducing 3 mL of 3% thioglycolate to mice via intraperitoneal injection. After 3 days, 10 mL of chilled PBS was introduced intraperitoneally to flush out macrophages. The cell suspension was then separated by centrifugation at 300 x *g* for 5 minutes. Cells were seeded in culture dishes containing DMEM with 10% FBS for 1 hour before being washed with serum-free DMEM two times to remove unattached cells. The cells were incubated in serum-free DMEM for 16 hours before treatment. To treat the macrophages, cells were incubated for 6 to 24 hours in DMEM without FBS with addition of LPS (Sigma-Aldrich), Pam3CSK4 (Invivogen), or SoL A. Cells were finally washed twice with Dulbecco’s phosphate-buffered saline before being lysed for total RNA or protein extraction.

### mRNA extraction and RT-qPCR in macrophage-based assays

Treated mouse macrophage cells were lysed with TriZol (Invitrogen) and total RNA was extracted from the cell lysate using a Direct-zol RNA miniprep kit (Zymo Research) according to the manufacturer’s protocol. The quality and quantity of RNA was then determined using a nanodrop and 1000 ng of mRNA from each sample was used for cDNA synthesis using a First-strand cDNA Synthesis System (Marligen Bioscience). qPCRs reactions were prepared in a 20 uL final volume containing Fast Start Universal SYBR Green Master (Rox) (Roche Applied Science), cDNA template, deionized water, and primers and probes for IL-1β, TNFα, IL-6, and the 18S rRNA which was used as a housekeeping gene Cycling conditions were 95 °C for 10 min followed by 40 cycles of 95 °C for 10 seconds, 60 °C for 15 seconds, and 68 °C for 20 seconds, then a melting curve analysis from 60 °C to 95°C every 0.2 °C was obtained. Amplifications were performed on an Eppendorf Realplex Mastercycler (Eppendorf). Relative gene expression levels were calculated using the ΔΔ*C_T_* method and expression levels of 18S were used to normalize the results.

### Molecular docking

The crystal structure of the TLR4/MD-2 comlpex was retrieved from the Protein Data Bank (PDB ID: 3FXI)^46^ and prepared using AutoDock Tools^83^. Molecular structures of SoL A, SoL B, sulfatide, and lipid A were constructed, and energy minimized using Marvin version 21.17.0, ChemAxon (https://www.chemaxon.com). Models of SoL A, SoL B, sulfatide, and lipid A were also prepared using AutoDock Tools and docked against the TLR4/MD-2 complex using AutoDock Vina^84, 85^ in a 32x32x32 angstrom box surrounding the MD-2 monomer. Docking results were visualized using PyMol^86^.

### ELISA displacement assay

Solid-phase sandwich ELISA kits were purchased from Invitrogen. The ELISA experiments were performed according to the kit instructions, using 50 nM hMD-2 (Novus Biologicals), 1 ng/mL LPS-EB Biotin (Invivogen), and 0.1, 1.0, and 10 μM purified SoL A. SoL A was added to the assay 1 hour before, 1 hour after, or simultaneously with LPS-EB Biotin. Absorbance was measured at 450 nm using a BioTek microplate reader.

### Western blot

Macrophages were treated with 100 ng/mL of LPS and 5 or 20 μM SoL A for 30 minutes. Following treatment, all cellular protein was extracted using MPER lysis buffer (Thermo Scientific). Protein samples were loaded onto SDS-PAGE gels for separation, then transferred to nitrocellulose membranes (Amersham Biosciences). Primary antibodies and HRP-conjugated secondary antibodies (Cell Signaling Technology) were used to detect target proteins. Signal was detected using an ECL kit (Thermo Scientific).

### Macrophage polarization

THP-1 monocytes were maintained in RPMI1640 with 10% heat inactivated FBS, 1% Penicillin-Streptomycin-Ampotericin B, and 50 μM 2-mercaptoethanol prior to differentiation. The cells were differentiated into macrophages with 150 nM PMA for 48 hours. M1 polarization was induced by adding 20 ng/mL IFN-γ and 100 pg/mL LPS for 24 hours. M2 polarization was induced by adding 20 ng/mL IL-4 and 20 ng/mL IL-13 for 24 hours. In all tests, 10 μM SoL A was added at the same time as M1 and M2 differentiation agents. After 24 hours of treatment, total RNA was collected, and RT-qPCR was performed as described above.

## Acknowledgements

This work is partially funded by a National Institutes of Health (NIH) grant P20GM103641, a National Science Foundation EPSCoR Program OIA-1655740, a Hong Kong Research Grants Council ECS grant HKU27107320, and the Hong Kong Branch of Southern Marine Science and Engineering Guangdong Laboratory (Guangzhou) (SMSEGL20SC02). We acknowledge Emily Quinn, Dorathea Lee, and Andrew Campbell for assistance with SoLs purification.

## Author contributions

E.A.O., J.Z., Y.-X.L., and J.L. designed research; E.A.O., J.Z., Z.F., D.X., Z.Z, M.K.M, M.M., and Y.W. performed research. All authors analyzed data and discussed results. All authors participated in preparing the manuscript.

## References

1. Donia, M. S. & Fischbach, M. A. Small molecules from the human microbiota. Science 349, 1254766 (2015).

2. Quinn, R. A. et al. Global chemical effects of the microbiome include new bile-acid conjugations. Nature 579, 123–129 (2020).

3. Lavelle, A. & Sokol, H. Gut microbiota-derived metabolites as key actors in inflammatory bowel disease. Nat Rev Gastroenterol Hepatol 17, 223–237 (2020).

4. Skelly, A. N., Sato, Y., Kearney, S. & Honda, K. Mining the microbiota for microbial and metabolite-based immunotherapies. Nat Rev Immunol 19, 305–323 (2019).

5. Spanogiannopoulos, P., Bess, E. N., Carmody, R. N. & Turnbaugh, P. J. The microbial pharmacists within us: a metagenomic view of xenobiotic metabolism. Nat Rev Microbiol 14, 273–287 (2016).

6. Cao, Y. et al. Commensal microbiota from patients with inflammatory bowel disease produce genotoxic metabolites. Science 378, eabm3233 (2022).

7. Yao, L. et al. A biosynthetic pathway for the selective sulfonation of steroidal metabolites by human gut bacteria. Nat Microbiol 7, 1404–1418 (2022).

8. Haiser, H. J. & Turnbaugh, P. J. Developing a metagenomic view of xenobiotic metabolism. Pharmacological Research 69, 21–31 (2013).

9. Flint, H. J., Scott, K. P., Duncan, S. H., Louis, P. & Forano, E. Microbial degradation of complex carbohydrates in the gut. Gut Microbes 3, 289–306 (2012).

10. Dai, H. et al. Recent advances in gut microbiota-associated natural products: structures, bioactivities, and mechanisms. Nat. Prod. Rep. 10.1039.D2NP00075J (2023) doi:10.1039/D2NP00075J.

11. Fischbach, M. A. Microbiome: Focus on causation and mechanism. Cell 174, 785–790 (2018).

12. Chaudhari, S. N., McCurry, M. D. & Devlin, A. S. Chains of evidence from correlations to causal molecules in microbiome-linked diseases. Nat Chem Biol 17, 1046–1056 (2021).

13. Brown, E. M., Clardy, J. & Xavier, R. J. Gut microbiome lipid metabolism and its impact on host physiology. Cell Host & Microbe 31, 173–186 (2023).

14. Koh, A., De Vadder, F., Kovatcheva-Datchary, P. & Bäckhed, F. From dietary fiber to host physiology: Short-chain fatty acids as key bacterial metabolites. Cell 165, 1332–1345 (2016).

15. Chiurchiù, V., Leuti, A. & Maccarrone, M. Bioactive lipids and chronic inflammation: Managing the fire within. Frontiers in Immunology 9, 38 (2018).

16. Bae, M. et al. Akkermansia muciniphila phospholipid induces homeostatic immune responses. Nature 608,168–173 (2022).

17. Durack, J. & Lynch, S. V. The gut microbiome: Relationships with disease and opportunities for therapy. Journal of Experimental Medicine 216, 20–40 (2018).

18. Lloyd-Price, J. et al. Multi-omics of the gut microbial ecosystem in inflammatory bowel diseases. Nature 569, 655–662 (2019).

19. Dziarski, R., Park, S. Y., Kashyap, D. R., Dowd, S. E. & Gupta, D. Pglyrp-regulated gut microflora *Prevotella falsenii*, *Parabacteroides distasonis* and *Bacteroides eggerthii* enhance and *Alistipes finegoldii* attenuates colitis in mice. PLOS ONE 11, e0146162 (2016).

20. Lima, S. F. et al. Transferable immunoglobulin A–coated *Odoribacter splanchnicus* in responders to fecal microbiota transplantation for ulcerative colitis limits colonic inflammation. Gastroenterology 162, 166–178 (2022).

21. Walker, A. et al. Sulfonolipids as novel metabolite markers of *Alistipes* and *Odoribacter* affected by high-fat diets. Scientific Reports 7, 11047 (2017).

22. MacEyka, M. & Spiegel, S. Sphingolipid metabolites in inflammatory disease. Nature 510, 58–67 (2014).

23. Olsen, I. & Jantzen, E. Sphingolipids in bacteria and fungi. Anaerobe 7, 103–112 (2001).

24. Pitta, T. P., Leadbetter, E. R. & Godchaux, W. Increase of ornithine amino lipid content in a sulfonolipid-deficient mutant of *Cytophaga johnsonae*. J Bacteriol 171, 952–957 (1989).

25. Hou, L. et al. Identification and biosynthesis of pro-inflammatory sulfonolipids from an opportunistic pathogen *Chryseobacterium gleum*. ACS chemical biology 17, 1197–1206 (2022).

26. Pasternak, B. A. et al. Lipopolysaccharide exposure is linked to activation of the acute phase response and growth failure in pediatric Crohn’s disease and murine colitis: Inflammatory Bowel Diseases 16, 856–869 (2010).

27. Caradonna, L. et al. Enteric bacteria, lipopolysaccharides and related cytokines in inflammatory bowel disease: biological and clinical significance. Journal of Endotoxin Research 6, 205–214 (2000).

28. Pastor Rojo, O., et al. Serum lipopolysaccharide-binding protein in endotoxemic patients with inflammatory bowel disease. Inflamm Bowel Dis 13, 269–277 (2007).

29. Im, E., Riegler, F. M., Pothoulakis, C. & Rhee, S. H. Elevated lipopolysaccharide in the colon evokes intestinal inflammation, aggravated in immune modulator-impaired mice. American Journal of Physiology-Gastrointestinal and Liver Physiology 303, G490–G497 (2012).

30. Almeida, A. et al. A unified catalog of 204,938 reference genomes from the human gut microbiome. Nat Biotechnol 39, 105–114 (2021).

31. Liu, Y. et al. Identification and characterization of the biosynthetic pathway of the sulfonolipid capnine. Biochemistry 61, 2861–2869 (2022).

32. Vences-Guzmán, M. Á., et al. Identification of the *Flavobacterium johnsoniae* cysteate-fatty acyl transferase required for capnine synthesis and for efficient gliding motility. Environmental Microbiology 23, 2448–2460 (2021).

33. Radka, C. D., Miller, D. J., Frank, M. W. & Rock, C. O. Biochemical characterization of the first step in sulfonolipid biosynthesis in *Alistipes finegoldii*. The Journal of biological chemistry 298, 1–13 (2022).

34. Radka, C. D., Frank, M. W., Rock, C. O. & Yao, J. Fatty acid activation and utilization by *Alistipes finegoldii*, a representative Bacteroidetes resident of the human gut microbiome. Molecular Microbiology 113, 807–825 (2020).

35. Kamiyama, T. et al. Sulfobacins A and B, novel von Willebrand factor receptor antagonists: I. Production, isolation, characterization and biological activities. The Journal of Antibiotics 48, 924–928 (1995).

36. Schirmer, M. et al. Dynamics of metatranscription in the inflammatory bowel disease gut microbiome. Nat Microbiol 3, 337–346 (2018).

37. Franzosa, E. A. et al. Gut microbiome structure and metabolic activity in inflammatory bowel disease. Nat Microbiol 4, 293–305 (2019).

38. Hale, L. P., Gottfried, M. R. & Swidsinski, A. Piroxicam treatment of IL-10-deficient mice enhances colonic epithelial apoptosis and mucosal exposure to intestinal bacteria. Inflammatory Bowel Diseases 11, 1060–1069 (2005).

39. Berg, D. J. et al. Rapid development of colitis in NSAID-treated IL-10-deficient mice. Gastroenterology 123, 1527–1542 (2002).

40. Cario, E. Toll-like receptors in inflammatory bowel diseases: A decade later. Inflammatory Bowel Diseases 16, 1583–1597 (2010).

41. Nothias, L.-F. et al. Feature-based molecular networking in the GNPS analysis environment. Nat Methods 17, 905–908 (2020).

42. Akira, S. & Takeda, K. Toll-like receptor signalling. Nat Rev Immunol 4, 499–511 (2004).

43. Kawasaki, T. & Kawai, T. Toll-like receptor signaling pathways. Frontiers in Immunology 5, 1–8 (2014).

44. Shimazu, R. et al. MD-2, a molecule that confers lipopolysaccharide responsiveness on Toll-like receptor 4. Journal of Experimental Medicine 189, 1777–1782 (1999).

45. Miyake, K. Roles for accessory molecules in microbial recognition by Toll-like receptors. Journal of Endotoxin Research 12, 195–204 (2006).

46. Park, B. S. et al. The structural basis of lipopolysaccharide recognition by the TLR4–MD-2 complex. Nature 458, 1191–1195 (2009).

47. Su, L. et al. Sulfatides are endogenous ligands for the TLR4–MD-2 complex. Proceedings of the National Academy of Sciences of the United States of America 118, 1–12 (2021).

48. Luk, J. M., Kumar, A., Tsang, R. & Staunton, D. Biotinylated lipopolysaccharide binds to endotoxin receptor in endothelial and monocytic cells. Analytical Biochemistry 232, 217–224 (1995).

49. Mosser, D. M. & Edwards, J. P. Exploring the full spectrum of macrophage activation. Nat Rev Immunol 8, 958–969 (2008).

50. Zhang, Y. et al. ECM1 is an essential factor for the determination of M1 macrophage polarization in IBD in response to LPS stimulation. Proc. Natl. Acad. Sci. U.S.A. 117, 3083–3092 (2020).

51. Zhang, X. & Mosser, D. Macrophage activation by endogenous danger signals. J. Pathol. 214, 161–178 (2008).

52. Parker, B. J., Wearsch, P. A., Veloo, A. C. M. & Rodriguez-Palacios, A. The genus *Alistipes*: Gut bacteria with emerging implications to inflammation, cancer, and mental health. Frontiers in Immunology 11, 906 (2020).

53. Chang, P. V., Hao, L., Offermanns, S. & Medzhitov, R. The microbial metabolite butyrate regulates intestinal macrophage function via histone deacetylase inhibition. Proc. Natl. Acad. Sci. U.S.A. 111, 2247–2252 (2014).

54. Trapecar, M. et al. Gut-liver physiomimetics reveal paradoxical modulation of IBD-related inflammation by short-chain fatty acids. Cell Systems 10, 223–239.e9 (2020).

55. Bhattarai, Y. et al. Bacterially derived tryptamine increases mucus release by activating a host receptor in a mouse model of inflammatory bowel disease. iScience 23, 101798 (2020).

56. Dodd, D. et al. A gut bacterial pathway metabolizes aromatic amino acids into nine circulating metabolites. Nature 551, 648–652 (2017).

57. Paik, D. et al. Human gut bacteria produce ΤΗ17-modulating bile acid metabolites. Nature 603, 907–912 (2022).

58. Sato, Y. et al. Novel bile acid biosynthetic pathways are enriched in the microbiome of centenarians. Nature 599, 458–464 (2021).

59. Li, W. et al. A bacterial bile acid metabolite modulates Treg activity through the nuclear hormone receptor NR4A1. Cell Host & Microbe 29, 1366–1377.e9 (2021).

60. Rakoff-Nahoum, S. & Medzhitov, R. Toll-like receptors and cancer. Nat Rev Cancer 9, 57–63 (2009).

61. Moreira Lopes, T. C., Mosser, D. M. & Gonçalves, R. Macrophage polarization in intestinal inflammation and gut homeostasis. Inflamm. Res. 69, 1163–1172 (2020).

62. Bhattarai, Y., Muniz Pedrogo, D. A. & Kashyap, P. C. Irritable bowel syndrome: A gut microbiota-related disorder? American Journal of Physiology-Gastrointestinal and Liver Physiology 312, G52–G62 (2016).

63. Wang, J. et al. A metagenome-wide association study of gut microbiota in type 2 diabetes. Nature 490, 55– 60 (2012).

64. Thomann, A. K. et al. Depression and fatigue in active IBD from a microbiome perspective-a Bayesian approach to faecal metagenomics. BMC Med 20, 366 (2022).

65. Mars, R. A. T. et al. Longitudinal multi-omics reveals subset-specific mechanisms underlying irritable bowel syndrome. Cell 182, 1460–1473.e17 (2020).

66. Buchfink, B., Reuter, K. & Drost, H.-G. Sensitive protein alignments at tree-of-life scale using DIAMOND. Nat Methods 18, 366–368 (2021).

67. Mirdita, M., Steinegger, M., Breitwieser, F., Söding, J. & Levy Karin, E. Fast and sensitive taxonomic assignment to metagenomic contigs. Bioinformatics 37, 3029–3031 (2021).

68. Chaumeil, P.-A., Mussig, A. J., Hugenholtz, P. & Parks, D. H. GTDB-Tk v2: memory friendly classification with the genome taxonomy database. Bioinformatics 38, 5315–5316 (2022).

69. Letunic, I. & Bork, P. Interactive Tree Of Life (iTOL) v5: an online tool for phylogenetic tree display and annotation. Nucleic Acids Research 49, W293–W296 (2021).

70. Li, H. & Durbin, R. Fast and accurate short read alignment with Burrows–Wheeler transform. Bioinformatics 25, 1754–1760 (2009).

71. Liao, Y., Smyth, G. K. & Shi, W. featureCounts: an efficient general purpose program for assigning sequence reads to genomic features. Bioinformatics 30, 923–930 (2014).

72. R Core Team. R: A language and environment for statistical computing. https://www.R-project.org/ (2021).

73. Oksanen, J. et al. vegan: Community ecology package. https://CRAN.R-project.org/package=vegan (2020).

74. Van Rossum, G. & Drake, F. L. Python 3 reference manual. (CreateSpace, 2009).

75. McKinney, W. Data structures for statistical computing in python. Proceedings of the 9th python in science conference, 51–56 (2010).

76. Virtanen, P. et al. SciPy 1.0: Fundamental algorithms for scientific computing in python. Nature Methods 17, 261–272 (2020).

77. Benjamini, Y. & Hochberg, Y. Controlling the false discovery rate: A practical and powerful approach to multiple testing. Journal of the Royal Statistical Society: Series B (Methodological) 57, 289–300 (1995).

78. Erben, U. et al. A guide to histomorphological evaluation of intestinal inflammation in mouse models. Int J Clin Exp Pathol 7, 4557–4576 (2014).

79. Ellermann, M. et al. Endocannabinoids Inhibit the Induction of Virulence in Enteric Pathogens. Cell 183, 650–665.e15 (2020).

80. Schmid, R. et al. Integrative analysis of multimodal mass spectrometry data in MZmine 3. Nat Biotechnol 41, 447–449 (2023).

81. Nothias, L.-F. et al. Bioactivity-based molecular networking for the discovery of drug leads in natural product bioassay-guided fractionation. J. Nat. Prod. 81, 758–767 (2018).

82. Shannon, P. et al. Cytoscape: A software environment for integrated models of biomolecular interaction networks. Genome Res. 13, 2498–2504 (2003).

83. Morris, G. M. et al. AutoDock4 and AutoDockTools4: Automated docking with selective receptor flexibility. J Comput Chem 30, 2785–2791 (2009).

84. Trott, O. & Olson, A. J. AutoDock Vina: Improving the speed and accuracy of docking with a new scoring function, efficient optimization, and multithreading. Journal of Computational Chemistry 31, 455–461 (2010).

85. Eberhardt, J., Santos-Martins, D., Tillack, A. F. & Forli, S. AutoDock Vina 1.2.0: New docking methods, expanded force field, and python bindings. J. Chem. Inf. Model. 61, 3891–3898 (2021).

86. Schrödinger, LLC. The PyMOL molecular graphics system, version 2.0. (2015).

